# Autoinflammatory patients with Golgi-trapped CDC42 exhibit intracellular trafficking defects leading to STING hyperactivation

**DOI:** 10.1101/2024.01.31.578189

**Authors:** Alberto Iannuzzo, Selket Delafontaine, Rana El Masri, Rachida Tacine, Giusi Prencipe, Masahiko Nishitani-Isa, Rogier T.A. van Wijck, Farzana Bhuyan, Adriana A. de Jesus Rasheed, Simona Coppola, Paul L.A. van Daele, Antonella Insalaco, Raphaela Goldbach-Mansky, Takahiro Yasumi, Marco Tartaglia, Isabelle Meyts, Jérôme Delon

## Abstract

Most autoinflammatory diseases are caused by mutations in innate immunity genes. Recently, four variants in the RHO GTPase CDC42 were discovered in patients affected by syndromes generally characterized by neonatal-onset of cytopenia and auto-inflammation, including hemophagocytic lymphohistiocytosis and rash in the most severe form (NOCARH syndrome). However, the mechanisms responsible for these phenotypes remain largely elusive. Here, we show that the recurrent p.R186C CDC42 variant, which is trapped in the Golgi apparatus, elicits a block in both anterograde and retrograde transports, and endoplasmic reticulum stress. Consequently, it favors STING accumulation in the Golgi in a COPI-dependent manner. This is also observed for the other Golgi-trapped p.*192C*24 CDC42 variant, but not for the p.Y64C and p.C188Y variants that do not accumulate in the Golgi. We demonstrate that the two Golgi-trapped CDC42 variants are the only ones that exhibit overactivation of the STING pathway. Consistent with these results, patients carrying Golgi-trapped CDC42 mutants present very high levels of circulating IFNα at the onset of their disease. Thus, we report new mechanistic insights on the impact of the Golgi-trapped CDC42 variants. This increase in STING activation provides a rationale for combination treatments for these severe cases.

## INTRODUCTION

Autoinflammatory diseases (AIDs) are due to genetic causes that drive hyperactivation of the innate immune system. They include clinically distinct disorders that present with systemic sterile inflammation, urticaria-like rashes, and disease-specific organ inflammation and damage. Some AIDs present with adaptive immune dysregulation, immunodeficiencies or with features of hyperinflammation, the latter including cytopenias, hyperferritinemia, hepatosplenomegaly, and high IL-18 levels (1–3). Most AIDs are caused by germline mutations in genes that encode key regulators of innate immunity.

We and others have previously reported heterozygous mutations in CDC42, encoding for a RHO GTPase which controls various cellular functions, including proliferation, migration, polarization, adhesion, cytoskeletal modifications and transcriptional regulation (4). Three types of CDC42 variants in the plasma membrane anchoring C-terminal region of CDC42 (c.556C>T, p.R186C; c.563G>A, p.C188Y and c.576A>C, p.*192C*24) have been found in patients with neonatal onset of AIDs (5–12). These patients show some common features such as immune deficiency, facial dysmorphisms, fever, polymorphic skin rashes, hepatosplenomegaly, neonatal-onset severe pancytopenia or dyshematopoiesis. Other unique features include macrothrombocytopenia and recurrent hemophagocytic lymphohistiocytosis (HLH) episodes in the most frequent and severe form of AID caused by the R186C substitution (NOCARH syndrome). Other variants affecting the same gene have been reported to variably affect CDC42 functional behavior and cause a clinically heterogeneous syndromic developmental disorder (13). Among these, a missense change (c.191A>G, p.Y64C) located in the N-terminal part of CDC42 underlies the Takenouchi-Kosaki syndrome, which is characterized by intellectual delay, growth retardation and dysmorphic facial features, macrothrombocytopenia and recurrent infections (14, 15). This peculiar phenotype has been also reported in a patient with a late onset systemic inflammatory disease and myelofibrosis (16). Although this patient presented with elevated levels of pro-inflammatory cytokines (IL-6, IL-18, IL-18BPa, CXCL9) as well as C-reactive protein (CRP) and erythrocyte sedimentation rate (ESR) compared to healthy controls, these levels were lower to those observed in NOCARH patients, suggesting a milder inflammation.

Thus, a better mechanistic characterization is essential for improved therapeutic management of these life-threatening conditions. In this study, we therefore sought to explore the mechanisms by which the CDC42 variants trigger severe AID. We show that the two disease-causing variants, R186C and *192C*24 which are trapped in the Golgi apparatus may exacerbate disease through STING activation and Type-I IFN-mediated pathology.

## RESULTS

Here, we report the further mechanistic characterization of seven unrelated CDC42 patients. Four patients (P1-4) carry the recurrent R186C variant present in the juxta-membranous polybasic region (5, 8, 10); two patients (P5-6) carry the *192C*24 variant due to a mutation in the stop codon which adds up a stretch of 24 amino acids at the C-terminal part of CDC42 (9, 10), while one patient (P7) carries the Y64C mutant located in the highly mobile Switch II region of CDC42 (16) (Figure 1A, B). These variants were systematically compared with the other C-terminal C188Y mutant which affects the cysteine residue of the CAAX sequence that is geranyl-geranylated and consequently, this variant cannot be lipidated (10).

**Figure 1:**
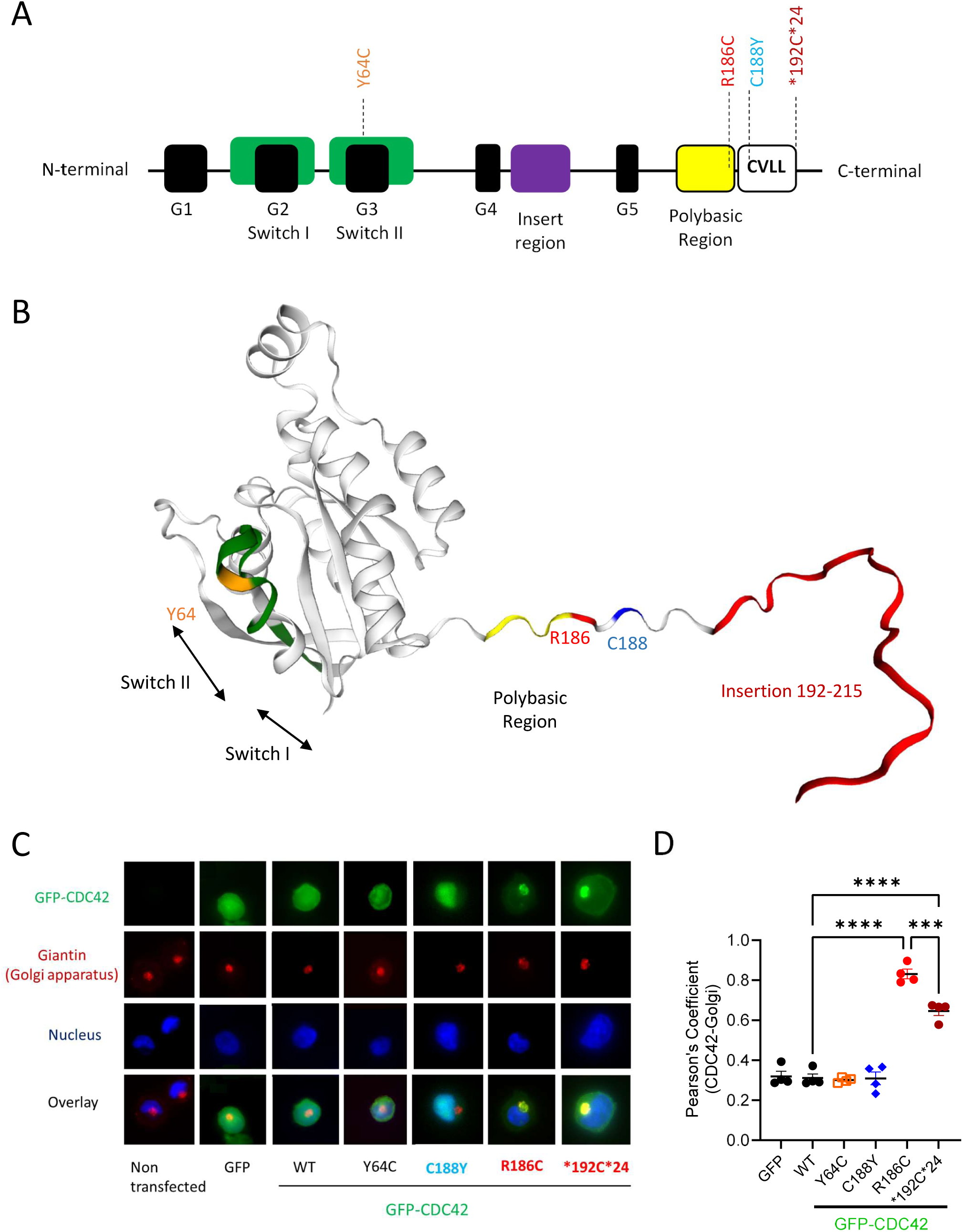
Localization of CDC42 variants identified in patients. **A,** Organization of CDC42 domains and indication of the positions of the pathogenic variants under study. **B,** Three-dimensional structure of CDC42 showing the CDC42 mutations of amino acids Y64, R186 and C188, and the insertion of the 24 amino acids at the C terminus highlighted in orange, red, blue, and dark red, respectively. The switch regions are indicated by black arrows and green color, and the polybasic region is represented in yellow. **C,** Subcellular localization of GFP-CDC42 variants in THP-1 cells co-stained for the Golgi and nucleus. **D,** Quantification of the degree of Golgi – CDC42 co-localization for different variants using the Pearson’s Coefficient. One dot represents the mean value from about 15 randomly-imaged cells from one independent experiment.

### CDC42 mutants exhibit a diverse subcellular localization

First, we compared the subcellular distribution of all these CDC42 variants in the THP-1 monocytic cell line using co-stainings with Golgi and nucleus markers (Figure 1C). The degree of colocalization between CDC42 and the Golgi was quantified using the Pearson’s Coefficient (Figure 1D), as described (11). We show here that the Y64C variant exhibits a normal diffuse subcellular localization. The non-membrane anchored C188Y mutant accumulates in the nucleus. In accordance with previous reports, the R186C and *192C*24 variants are the only ones to exhibit a retention in the Golgi due to aberrant palmitoylation (5, 11, 17). Moreover, we demonstrate that the R186C variant is quantitatively trapped more in the Golgi compared to the *192C*24 mutant (Figure 1D).

### CDC42 R186C induces a block in anterograde trafficking

Because of the peculiar retention of two C-terminal CDC42 mutants in the Golgi, we next wondered whether they affect the Golgi function, and in particular the intracellular bidirectional trafficking between the endoplasmic reticulum (ER) and the Golgi.

To study the anterograde ER-to-Golgi transport, we followed the trafficking of endogenous proto-collagen of type I (PC-1) as a protein model (18). For this purpose, fibroblasts from four healthy donors (HD) or from Y64C and R186C patients were co-stained for Golgi, ER and nucleus markers together with PC-1 (Figure 2A). The initial high temperature condition of cell culture is necessary for accumulating the majority of PC-1 in the ER at time 0 as shown by the quantification of the Pearson’s Coefficient of 0.6-0.7 for all conditions (Figure 2B). The accumulation of PC-1 is initially low in the Golgi as shown by a low Pearson’s Coefficient of about 0.4 in all cells (Figure 2C). Upon decreasing the temperature to 32°C, PC-1 proteins are released from the ER and are expected to follow the normal secretory route through the Golgi apparatus. Quantifications of the Pearson’s Coefficients after a 1-hour shift in temperature show that cells from all four HD and from the Y64C patient indeed exhibit a drop in PC-1 - ER colocalization to 0.5 and an increase in PC-1 - Golgi colocalization to 0.6-0.7 (Figure 2B, C). By contrast, cells endogenously expressing the CDC42 R186C mutant exhibit Pearson’s Coefficients that remain high for the PC-1 - ER colocalization and low for the PC-1 - Golgi colocalization, similar to the ones observed at basal levels (time 0). This indicates that the anterograde trafficking is severely impaired in R186C patient cells.

**Figure 2:**
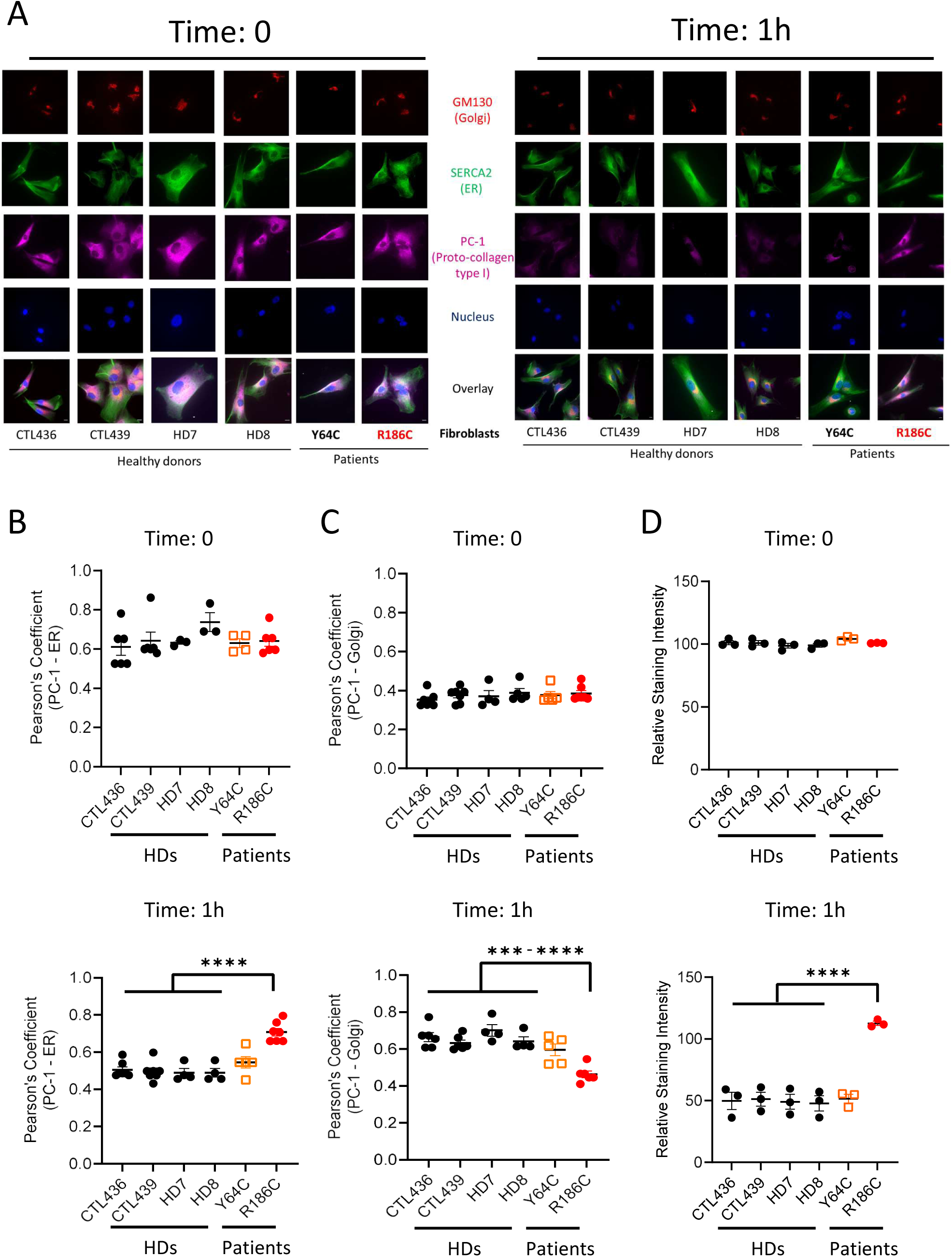
Impact of CDC42 variants on anterograde transport. **A**, Immunofluorescence analyses of PC-1, ER, Golgi and nucleus localizations in 4 different healthy donor (HD) fibroblasts, patients Y64C and R186C at 0 and 1h. Images are representative of at least 3 independent experiments. Quantifications of the colocalizations between PC-1 and the ER (**B**) or the Golgi (**C**) at the indicated times for each cell type. **D,** Analysis of total PC-1 fluorescence intensity following anterograde transport. The relative intensity of PC-1 expression was normalized to 100 from the mean values of the 4 healthy donors at time 0. In all graphs, one dot represents the mean value from about 15 randomly-imaged cells from one independent experiment.

We also quantified the total amount of PC-1 expression in all these fibroblasts. Initially, all cells express similar total levels of PC-1 (Figure 2D). However, upon 1 hour of temperature shift, all HD and the Y64C fibroblasts show a 50% reduction in the expression levels of PC-1, probably due to the processing and secretion of a fraction of PC-1 proteins. In contrast, PC-1 expression levels slightly increased in fibroblasts from the CDC42 R186C patient. Altogether, these results indicate that the Golgi-trapped CDC42 R186C variant exhibits a specific block of the anterograde ER - Golgi trafficking.

### CDC42 R186C inhibits retrograde transport

To investigate the functionality of the retrograde Golgi - ER transport in these same cells, we used the Cholera toxin B subunit (CtxB) model (19) with costainings for Golgi, ER and nucleus (Figure 3A). Upon internalization, most CtxB reaches the Golgi. For all fibroblasts, the CtxB-Golgi colocalization was initially similar (Figure 3B) and higher than the CtxB-ER colocalization (Figure 3C) (Pearson’s Coefficient of 0.7 *versus* 0.5). Eight hours later, the reverse was true in HD and Y64C cells, as most CtxB had reached the ER (Pearson’s Coefficient: 0.7) and fewer were still present in the Golgi (Pearson’s Coefficient: 0.5). Conversely, no changes were observed for the R186C patient cells indicating that the Golgi-trapped CDC42 R186C variant also impairs the retrograde Golgi - ER trafficking pathway.

**Figure 3:**
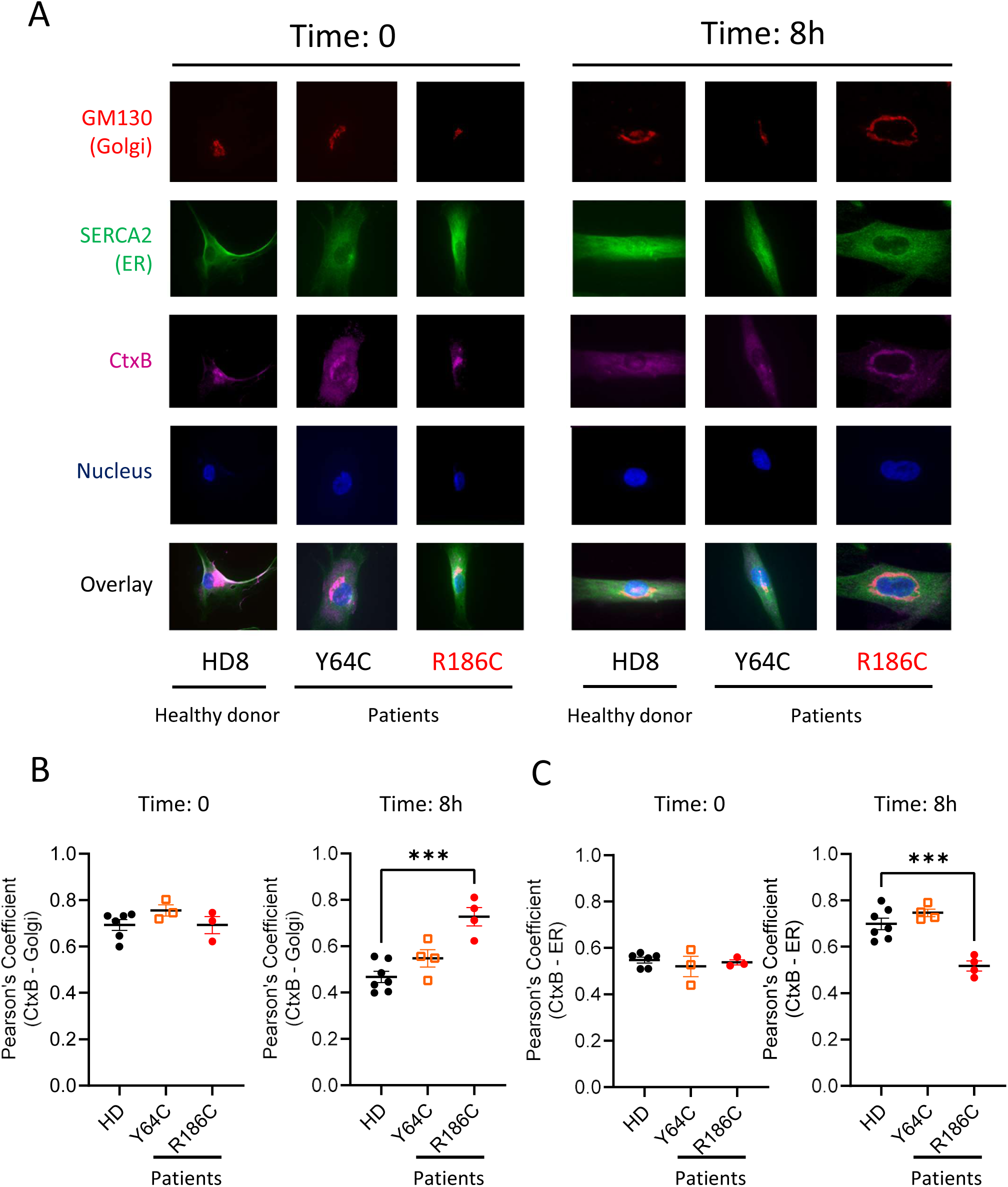
Impact of CDC42 variants on retrograde transport. **A**, Immunofluorescence analyses of retrograde transport assay of Cholera toxin B subunit (CtxB) in fibroblasts from the HD8 healthy donor and from the Y64C and R186C CDC42 patients at times 0 and 8 h. Co-stainings for Golgi, ER and nuclei were performed. Images are representative of at least 3 independent experiments. Quantifications of microscopy images are shown for the CtxB - Golgi (**B**) and CtxB - ER (**C**) colocalizations at the indicated times. Transport assay data from 4 healthy donors were pooled. In all graphs, one dot represents the mean value from about 15 randomly-imaged cells from one independent experiment.

### CDC42 R186C triggers ER stress

Next, we hypothesized that the defective intracellular trafficking caused by the CDC42 R186C pathogenic mutant could increase ER stress. Accordingly, we found an increased expression of the molecular chaperone binding immunoglobulin protein (BiP) by microscopy in baseline conditions in CDC42 R186C patient’s cells but not in HD and Y64C cells (Figure 4). Thus, the impairments in both anterograde and retrograde trafficking observed in cells heterozygous for the CDC42 R186C mutation are likely to be responsible for ER stress which has been shown to be related to inflammation (20).

**Figure 4:**
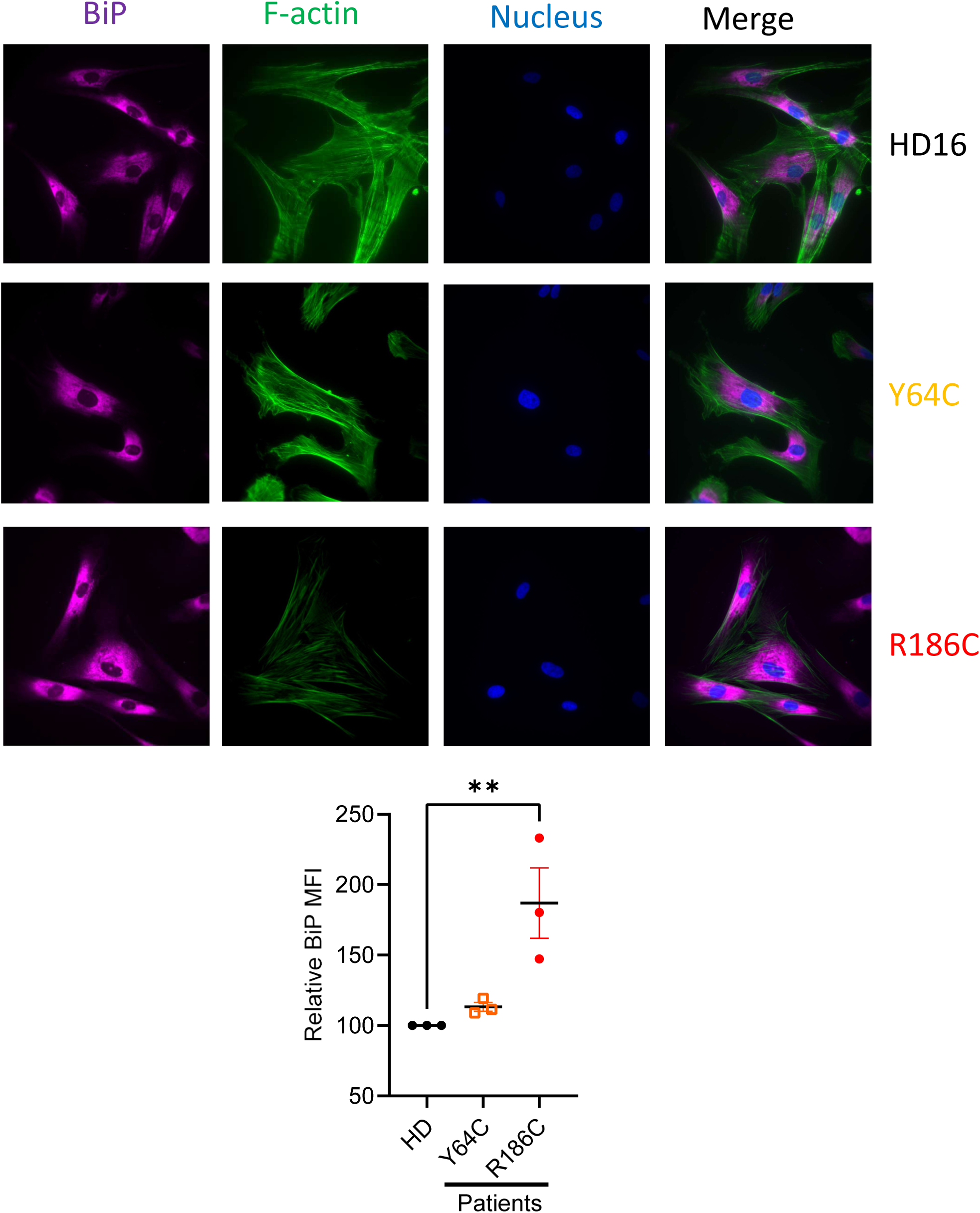
Impact of CDC42 variants on ER stress: Immunofluorescence analyses of ER stress in fibroblasts from the HD16 healthy donor and from the Y64C and R186C CDC42 patients. Co-stainings for BiP protein, F-actin and nuclei were performed. Images are representative of at least 3 independent experiments. Quantification of microscopy images is shown below. In all graphs, one dot represents the mean value from about 15 randomly-imaged cells from one independent experiment.

### Golgi-trapped CDC42 mutants induce STING hyperactivation

As previously described, the small pool of CDC42 naturally present in the Golgi apparatus in physiological conditions can interact with the COPI complex and regulate COPI-dependent bidirectional transport (21–23). In this report, we have shown that an excess of pathogenic CDC42 in the Golgi blocks the bidirectional ER-Golgi route. Interestingly, we and others have shown that this particular defect is also observed in patients with heterozygous mutations in *COPA* (24–27), encoding for one of the subunits of the COPI complex. As these COPA patients have been shown to develop inflammatory syndromes because of alterations in STING trafficking, we next studied whether Golgi-trapped CDC42 variants also exhibit changes in the subcellular localization of STING.

To this aim, we stained patients’ and HD cells for STING, Golgi and nucleus. We observed STING enrichment in the Golgi from the R186C patient cells but not in Y64C or HD cells (Figure 5A). Furthermore, in accordance with this, ectopic overexpression of WT, Y64C and C188Y CDC42 in THP-1 cells did not impact STING localization (Figure 5B). By contrast, the two Golgi-trapped variants R186C and *192C*24 elicited STING accumulation in the Golgi. This enrichment of STING was abrogated when these mutants were precluded from interacting with the COPγ subunit of COPI by mutating the K183K184 motif in CDC42 (21) (Figure 5C). These data point to the crucial interaction of CDC42 with COPI for intracellular transport. Thus, although the CDC42 K183S/K184S/R186C and K183S/K184S/*192C*24 mutants retain a strong Golgi accumulation (Figure 5D), only CDC42 R186C and *192C*24 trigger STING accumulation in the Golgi, and this effect depends on the interaction of CDC42 with COPI. Because STING is only phosphorylated and active when it reaches the Golgi (28–30), we next monitored several downstream readouts / effectors of the STING pathway. Phosphorylation of IRF3 (P-IRF3) was specifically increased only in THP-1 cells expressing the R186C and *192C*24 variants (Figure 5E). R186C patient cells also showed higher levels of P-IRF3 (Figure 5F) and P-STAT1 (Figure 5G). Altogether, these results indicate that the STING signaling pathway is hyperactivated in cells expressing the Golgi-trapped CDC42 variants.

**Figure 5:**
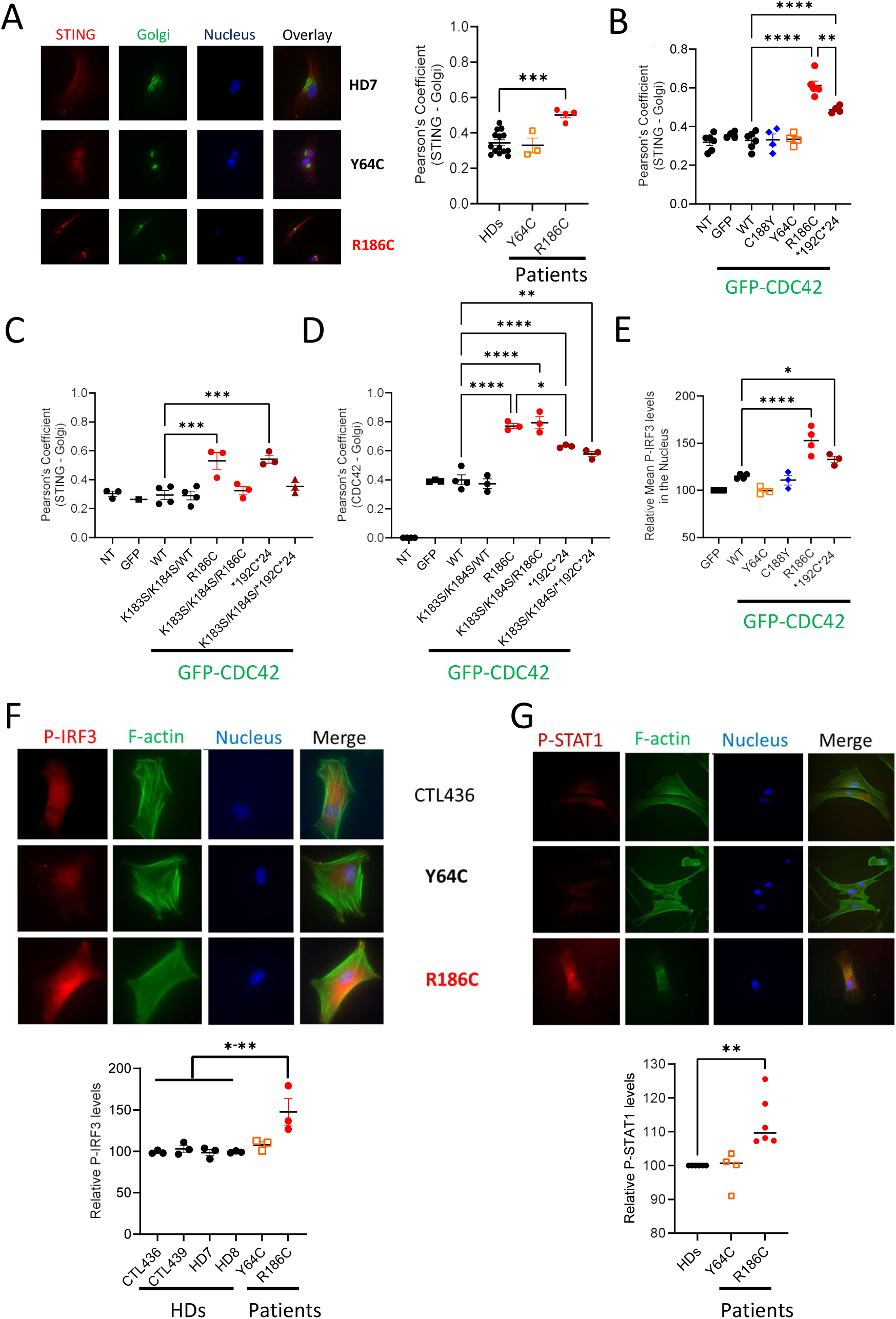
Golgi-trapped CDC42 variants induce STING accumulation in the Golgi and activation. **A**, Left: Subcellular stainings of STING, Golgi and nuclei in controls and patients’ fibroblasts. Right: Quantification of the degree of STING - Golgi co-localization in each condition. 4 controls were pooled. **B,** Measurement of STING - Golgi co-localization in THP-1 cells expressing different forms of CDC42. Quantifications of the degrees of STING - Golgi (**C**) or CDC42 - Golgi (**D**) co-localizations in THP-1 cells expressing different mutants of CDC42, including some with the K183S/K184S double mutation which inhibits COPI binding. NT: non transfected. **E,** Measurement of P-IRF3 intensity in the nuclei of THP-1 cells expressing GFP-CDC42 variants after normalization to 100 for THP-1 cells expressing GFP alone. Immunocytochemistry stainings and quantifications of P-IRF3 (**F**) and P-STAT1 (**G**) intensities in healthy donors or CDC42 patients’ fibroblasts. Results were normalized by setting the value 100 for controls. In all graphs, one dot represents the mean value from about 15 randomly-imaged cells from one independent experiment.

### Golgi-trapped CDC42 patients show increased levels of IFNα at the onset of the disease

To test the possibility that the hyperactivation of the STING pathway elicits increased type I IFN (IFN-I), we first measured the levels of IFNα present in available serum or plasma samples from Golgi-trapped CDC42 patients at different stages (onset, prior to bone marrow transplantation, late) and under different treatments (Figure 6; Supplemental material; Supplemental Table 1). R186C patients P1-4 all had very high levels of IFNα at the onset of the disease or after early treatment. These amounts of IFNα were reduced to various extent upon diverse treatments or upon bone marrow transplantation, although basal levels as in HD were not systematically reached. *192C*24 Patient 5 also had very high levels of IFNα at an early time point which was resolved later on. *192C*24 Patient 6 was on treatment with various doses of corticosteroids during her life due to recurrent vasculitis. This may explain why her IFNα levels were systematically low in the measured samples (Figure 6).

**Figure 6:**
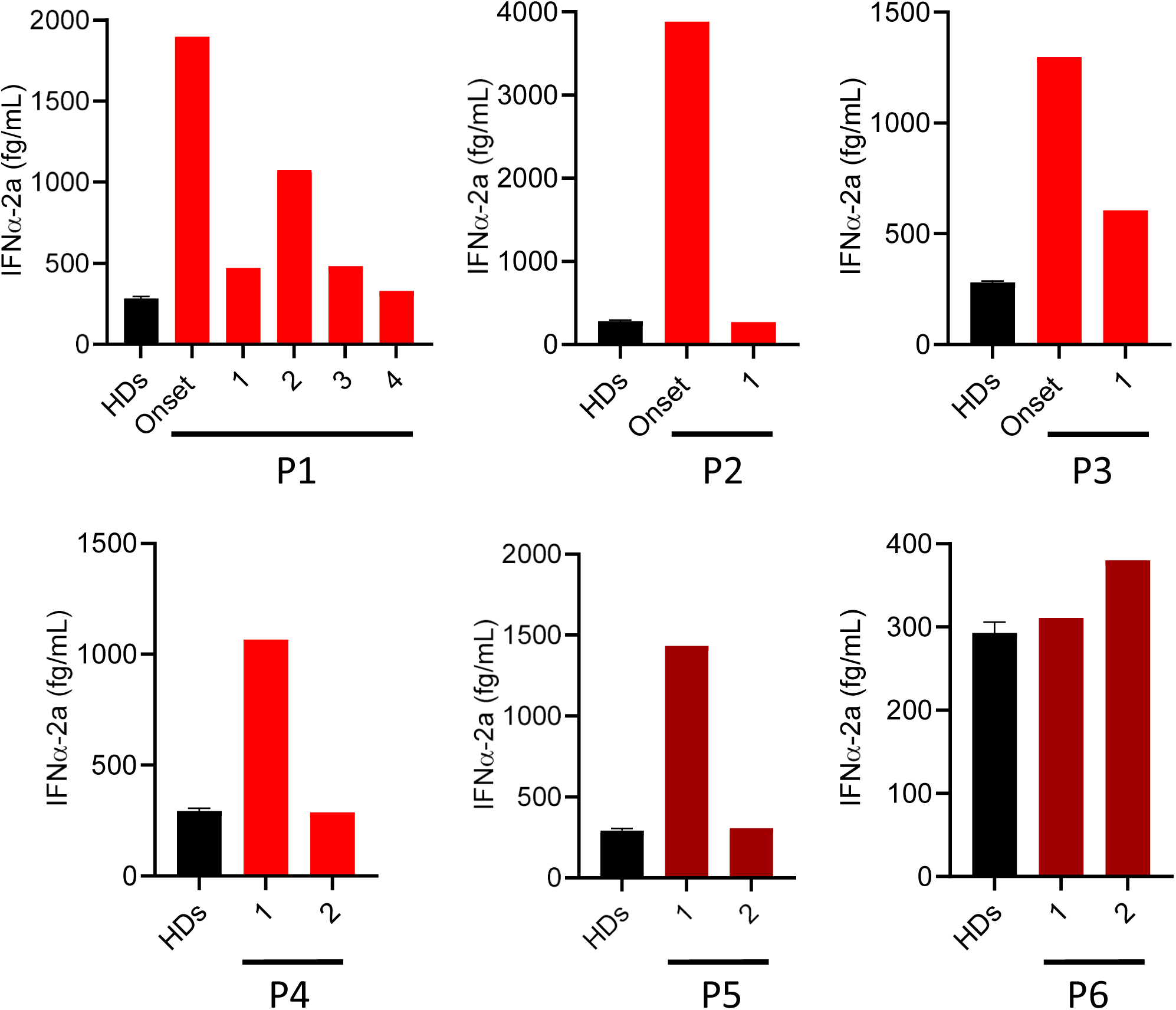
Golgi-trapped CDC42 variants are associated with high IFNa levels in the blood. Quantification of IFNa in sera or plasma from R186C (P1-4) and *192C*24 (P5-6) CDC42 patients at the onset of the disease or after different treatments: P1 → 1: probable virosis (ongoing glucocorticoids and anakinra), 2: infection by adenovirus and rhinovirus (ongoing glucocorticoids, anakinra and cyclosporine), 3: ongoing cyclosporine, glucocorticoids and anakinra, 4: bone marrow transplantation. P2 → 1: high dose of glucocorticoids and anakinra. P3 → Onset: improved state. 1: Re-inflammatory state, anti-TNF treatment. P4 → 1: anakinra, 2: glucocorticoids and canakinumab. P5 → 1: anakinra, 2: high dose of anakinra. P6 → 1 and 2: corticosteroids and intravenous Ig (Immunoglobulin response therapy).

### Golgi-trapped CDC42 mutants induce high expression of IFN-stimulated genes (ISG)

Next, we measured the mRNA expression of six IFN-stimulated genes (ISG) to calculate the IFN score. Contrary to Y64C patients’ cells, the R186C cells showed a marked increase in IFN score (Figure 7A). This was also confirmed in THP-1 cells where only the two Golgi-trapped variants showed a high IFN score whereas the Y64C and C188Y mutants failed to do so (Figure 7B). Furthermore, although the IFN score was slightly reduced in cGAS KO THP-1 cells (Figure 7C, D), R186C and *192C*24 expressions were still able to elicit higher ISG production compared to WT CDC42. However, the increased ISG expression in R186C and *192C*24 was dependent on STING as it was completely abrogated in STING KO THP- 1 cells (Figure 7E). This indicates a strict requirement for STING to induce ISG expression by these two Golgi-trapped CDC42 variants.

**Figure 7:**
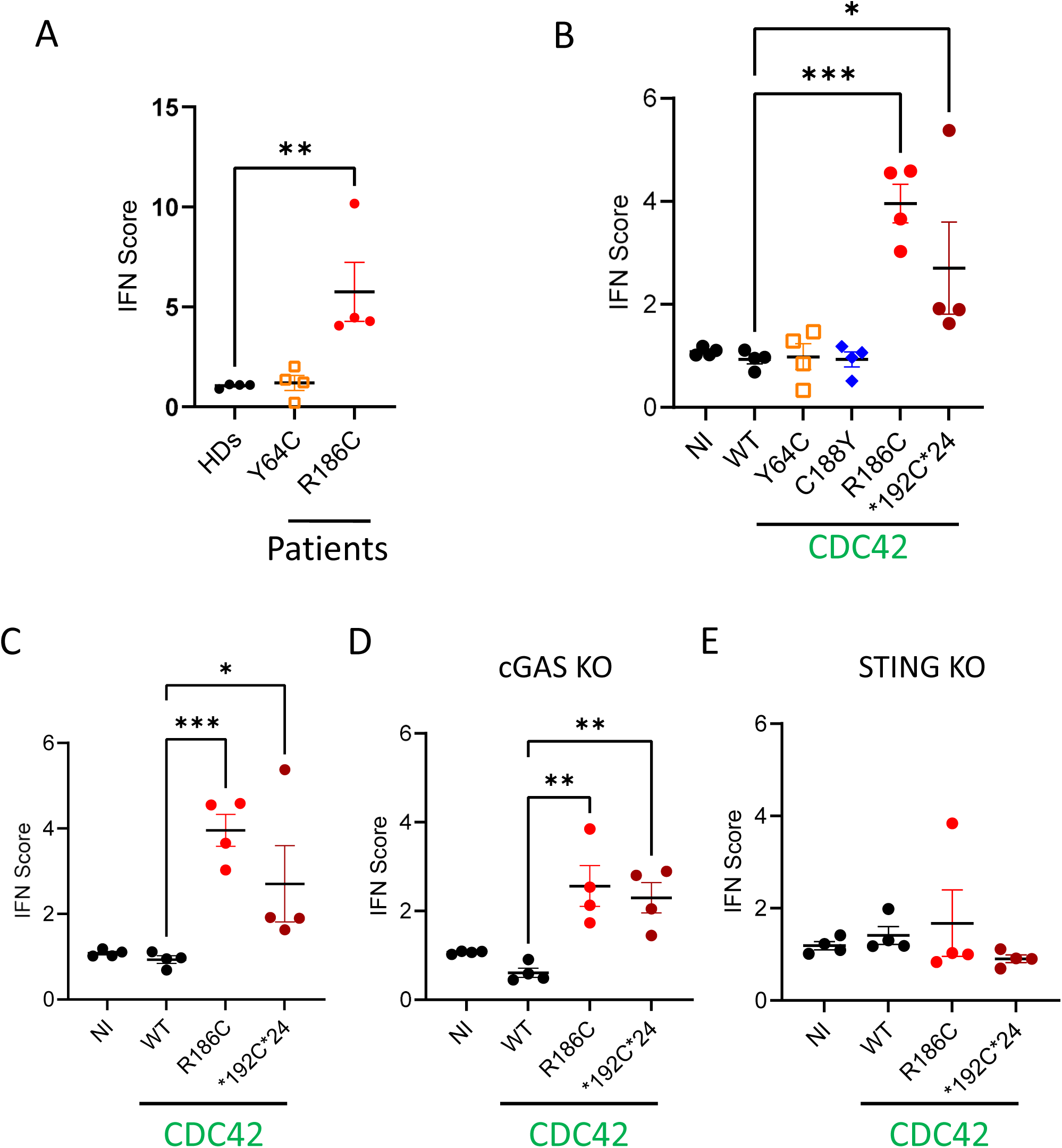
Golgi-trapped CDC42 variants induce increased Interferon signature gene expression in a STING-dependent way. IFN scores (median value of the 6 ISG: *IFI27*, *IFI44L*, *IFIT1*, *ISG15*, *RSAD2* and *SIGLEC1*) from 4 independent experiments were quantified in patients’ fibroblasts (**A**), THP-1 (**B**, **C**), THP-1 cGAS KO (**D**) and THP-1 STING KO (**E**) cells expressing different CDC42 mutants. Each dot represents the IFN score of one independent experiment. Panels B and C are partly duplicated. NI: non-infected.

## DISCUSSION

Overall, we extend the spectrum of dysfunctions previously observed in Golgi-trapped CDC42 patients (5–12). More specifically, we provide new and specific mechanistic insights for the CDC42 R186C and *192C*24 variants.

Indeed, we show that the recurrent CDC42 R186C mutant, reported now in 13 patients affected by autoinflammation, impairs anterograde and retrograde trafficking between the ER and the Golgi, and elicits ER stress. This in turn leads to STING accumulation in the Golgi as we and others previously reported for patients carrying mutations in genes involved in the transport machinery such as *COPA* (24–27) and, more recently, *ARF1* (31). Importantly, we demonstrate that the interaction between COPI and the Golgi-trapped CDC42 mutants is absolutely required for triggering STING enrichment in the Golgi. Of note, the R186C mutant is more efficient than the *192C*24 one, and this correlates with a higher trapping of R186C in the Golgi compared to the *192C*24 mutant. Consequently, only these two Golgi-localized CDC42 mutants induce STING activation and ISG expression. In our hands, the other C-terminal C188Y mutant, which is not trapped in the Golgi, does not induce STING accumulation in the Golgi, nor STING activation, nor ISG expression. Our data also suggest that ISG expression, induced by both Golgi-trapped CDC42 mutants, is largely cGAS-independent which seems at odds with the recent report of defective mitochondria releasing DNA in the cytosol of R186C and C188Y fibroblasts (12). Maybe CDC42 patient fibroblasts but not THP-1 expressing pathogenic CDC42 variants are the only cells showing an impairment of the mitochondria structures. Using STING-deficient cells, our data support a compulsory requirement for STING expression for inducing ISG expression specifically by Golgi-trapped CDC42 mutants. Thus, there seems to be several possible mechanisms to hyperactivate the STING pathway in these pathological conditions. Finally, the Y64C mutant does not exhibit any defects in the readouts we studied, probably in line with the late onset of the disease for this patient (16). Potentially, other mechanisms are at play in this variant.

ER stress plays a complex role in the induction of inflammation (20). Defects in the retrograde Golgi-to-ER transport caused by mutations in different COPI subunit proteins, were previously described to result in an increased ER stress (27,32–35). Similarly, we observed an increased ER stress specifically in CDC42 R186C. One can assume that the increased ER stress, that we have observed here, could result from an impaired retrieval of chaperone proteins to the ER, due to a strongly reduced Golgi-to-ER retrograde transport. In addition, accumulation of nascent proteins in the ER, due to impairment in ER-to-Golgi anterograde transport induced by CDC42 R186C, could also be responsible for ER stress. Increased ER stress together with defective bidirectional trafficking are therefore key pathogenic mechanisms caused by the CDC42 R186C variant responsible for the NOCARH syndrome.

Thus, next to increased NF-κB (11) and Pyrin (8, 36) activation, high ER stress and STING hyperactivation are here unveiled as additional proinflammatory pathways elicited by the pathogenic CDC42 Golgi-trapped variants in both patients’ cells and upon ectopic expression. In agreement with the hyperactivation of the STING pathway, we show marked increases in IFNα concentration for all five unrelated R186C and *192C*24 CDC42 patients for whom blood samples at the onset of the disease or upon early treatment were available. These high IFNα levels were reduced to normal upon transplantation, indicating that mutated immune cells are responsible for the observed defects.

In this way, our results may lead to novel insights in treatment approaches, by targeting the affected pathways in these patients. Moreover, our findings showcase a layer of complexity regarding the extreme variability in immunological and cellular phenotypes provoked by pathogenic variants in a single gene despite all patients being affected by autoinflammation.

## METHODS

### Study oversight

Human studies were carried out according to French law on biomedical research and to the principles outlined in the 1975 Helsinki Declaration and its modification. Institutional review board approvals were obtained (DC-2023-5921 and IE-2023-3004 from the French Ministry of Higher Education and Research). All patients provided written informed consent for the conservation and use of their blood samples and cells for research.

### *In silico* CDC42 structural analysis

The SWISS-MODEL server (https://swissmodel.expasy.org/) was used based on the template of the three-dimensional X-ray structure of CDC42 (PDB accession number: 5c2j.1.B). It shows the well-known structure of CDC42 and predicts the additional 24 amino to the protein’s C terminus present in the *192C*24 mutant. This additional sequence is predicted to be an unstructured region with a low confidence level.

### Plasmid constructs

The GFP-CDC42 plasmid previously described (37) encodes for the ubiquitous isoform 1 of CDC42 and contains a GFP tag in N-terminal. From this construct, R186C, K183S/K184S/R186C, C188Y and Y64C) were obtained by site-directed mutagenesis (Quick change kit, Agilent Technologies). The *192C*24 and K183S/K184S/*192C*24 plasmids were produced by ThermoFisher. The pmaxGFP vector was provided by Lonza. Plasmids encoding for Gag, Pol and VSV-G were provided by Thomas Henry (*Centre International de Recherche en Infectiologie*, Lyon, France). Nucleotide coding sequences of myc-tagged CDC42 WT and variants cloned into the pLenti-EF1a-IRES-EGFP (Creative Biolabs) were provided by the CIGEx facility (CEA, Fontenay-aux-Roses, France).

### Cells

Primary fibroblasts from the Y64C patient (P7) and healthy donors were obtained from the Leuven biobank of primary immunodeficiency. Primary fibroblasts from the R186C patient and healthy donors were previously described (11). They were cultured in DMEM medium supplemented with 10 % Fetal Calf Serum, antibiotics (Pennicilin and Streptomycin) and sodium pyruvate. The human THP-1 monocytic cell line was provided by Serge Bénichou (*Institut Cochin*, Paris). THP-1 cGAS KO and STING KO were kindly provided by Marie-Louise Frémond (*Institut Imagine*, Paris). They were cultured in RPMI medium supplemented with 10 % Fetal Calf Serum, antibiotics (Pennicilin and Streptomycin) and sodium pyruvate. All cells were regularly tested for Mycoplasma (Lonza).

### Patients Samples

For serum extraction, blood was collected in an SST tube, spun at 3,000 rcf for 10 minutes and stored at −20°C. For plasma extraction, blood was collected in EDTA tubes, spun twice at 2,000 rcf for 10 and 15 minutes. Sera and plasmas from healthy donors used as controls were collected from age and sex-matched individuals.

### Transfections

2×10^6^ THP-1 cells were centrifuged and washed in PBS (Gibco). They were then transfected by nucleofection with 3 μg of plasmid DNA in 100 μl of Cell Line Nucleofector Solution V (Lonza) using the V-001 program (Amaxa Biosystems). After transfection, 500 μL of warmed and complete RPMI medium was added to the cells, which were then plated into 6-well plates containing 1.5 mL of the same medium. The plates were then incubated overnight.

### Lentiviral transduction

5×10^6^ HEK-293T cells were seeded in 10 cm-diameter Petri dishes in complete DMEM and left overnight at 37°C with 5% CO2. 2h before the transfection, the medium was replaced and the following mix was prepared: Plasmid of interest (20 μg), pPAX 8.91 gag-pol expressor (10 μg), pMDG VSV-G expressor (5 μg), 1 M CaCl_2_ and H_2_O (qsp 0.5 mL). This mix was added to 0.5 mL of 2X HBS, incubated at room temperature for 30 min and added overnight to the dishes. The next day, the medium was removed and replaced with 5 mL of DMEM. 24 h later, the media were recovered, centrifuged at 1,000 g for 5 min, and filtered with a 0.45 µm filter. After addition of a sucrose solution (200 g/L sucrose, 100 mM NaCl, 20 mM HEPES, 1 mM EDTA), the samples were centrifuged for 2 h at 21.109 rpm (82.700 g) at 4°C. The supernatant was removed and the pellet was resuspended in 100 μL of PBS 1X without Ca^2+^ and Mg^2+^ and incubated at 4°C for 2h. Lentiviruses were then stored at −80°C. 8×10^4^ THP-1 cells were infected with viruses, and analyzed by flow cytometry (BD FACSCalibur) at day 4 after infection. Infected cells were identified by GFP expression and sorted by flow cytometry (FACS ARIA III, BD biosciences).

### Trafficking assays

These assays were performed as described (27). Briefly, fibroblasts were first seeded on labtechs (Chamber SlideTM system 154534 - ThermoFisher).

For testing the anterograde transport, we favored PC-1 retention in the ER by incubating the cells for 3h at 40°C in complete DMEM (18). Cells were then shifted to 32°C for 1h.

For the evaluation of the retrograde transport, cells were incubated with 0.15 μg/mL AF555-labeled CtxB (Molecular Probes, C34776) on ice for 30 minutes (19). After washing twice with PBS, the cells were incubated with a pre-warmed DMEM medium for the indicated times at 37°C. Time 0 corresponds to the maximal retention of CtxB in the Golgi and is obtained after 2 hours of incubation. CtxB maximally reaches the ER 8 hours later.

### ER stress assay

To evaluate the ER stress, fibroblasts were fixed and permeabilized as described previously (27). They were then incubated for 45 min with anti-GRP78 BiP antibody, and F-actin was labeled with phalloidin AlexaFluor488. Cells were then washed twice with 1.5 mL of saponin buffer to remove non-specifically bound antibodies and incubated with a secondary antibody for 30 minutes in the dark. Cells were washed twice in saponin buffer and once with 500 μL of PBS. The staining of the nuclei was next performed with 1 μg/mL Hoechst for 10 min in the dark. Cells were finally washed twice with 1.5 mL PBS and MilliqWater.

### Intracellular fluorescent stainings

Cells were fixed with 4 % paraformaldehyde (Electron Microscopy Sciences) for 10 min for all experiments except for P-STAT1 staining that used a cold 90 % - methanol solution in ice for 15 min. Then, cells were washed once in saturation buffer [PBS 1 % BSA (Sigma)], permeabilized with 0.1% Triton buffer for 10 minutes and washed twice in permeabilization buffer (PBS containing 0.1 % saponin (Fluka Biochemika) and 0.2 % BSA). Cells were then incubated for 45 min with the primary antibodies based on the type of experiment (Supplemental Table 3). They were then washed twice with 1.5 mL of saponin buffer and incubated with secondary antibodies (Supplementary Table 3) for 30 min in the dark. Cells were washed twice in saponin buffer and twice with PBS. Nuclei labeling was next performed with 1 µg/mL of Hoechst for 10 min in the dark. Cells were finally washed twice with 1.5 mL PBS, and mounted with VECTASHIELD (Vector Laboratories H-1700).

### Meso Scale Discovery multiplex assay

S-PLEX human Interferon KIT (K151P3S-1, for IFNα-2a) was purchased from Mesoscale Discovery (MSD). S-PLEX plates were coated with linkers and biotinylated capture antibodies, according to manufacturer’s instructions. The assay was performed according to the manufacturer’s protocol with overnight incubation of the diluted samples and standards at 4°C. The electrochemiluminescence signal (ECL) was detected by MESO QuickPlex SQ 120 plate reader and analyzed with Discovery Workbench Software (v4.0, MSD). The concentration of each sample was calculated based on the parameter logistic fitting model generated with the standards. The concentration was determined according to the certificate of analysis provided by MSD.

### RNA isolation and qRT-PCR

For RNA extraction, we used the PureLink RNA mini Kit (Invitrogen 12183018A) according to the manufacturer’s instructions. mRNA were reverse-transcribed with Superscript Vilo cDNA synthesis kit (Thermo Fisher Scientific, 11754-050). qPCR was performed using SYBR green (Bio-rad, 1725271). The CT values obtained for the genes of interest were corrected for the cDNA input by normalization to the CT value of GAPDH (ΔCT). Furthermore, the ΔCT value was normalized to the mean of control non-transduced cells. Finally, by using the formula 2−ΔΔCT, the relative quantification of target cDNA was described as a fold-increase above control and normalized to GAPDH. A list of the probes used is supplied in Supplementary Table 2. For each independent experiment, we averaged the triplicate values and this mean was used to calculate the IFN score as the median values of 6 ISG (*IFI27*, *IFI44L*, *IFIT1*, *ISG15*, *RSAD2* and *SIGLEC1*).

### Imaging

Images were obtained with an inverted fluorescence microscope (Eclipse Nikon TE300), a Photometrics Cascade camera, and acquired using the Metamorph v7.8.9.0 software. A 100X or 40X objective was used for THP-1 cells and fibroblasts, respectively.

Images analyses were performed using Fiji software (ImageJ version 1.51u). The Pearson’s Coefficient was measured in Fiji by using a macro containing the Coloco2 plugin. This coefficient measures the degree of overlap between two stainings. A Pearson’s Coefficient value of 0 means that there is no colocalization between the two stainings. By contrast, a value of 1 indicates that there is a perfect colocalization between the two stainings under study. Fibroblasts staining intensity was measured in the whole cells by taking into account the cell morphology identified by F-actin phalloidin staining. P-IRF3 staining intensity in THP-1 cells was measured by focusing and limiting the quantification of the staining to the nucleus. Each condition was normalized to the intensity values measured in healthy donor cells or THP-1 cells expressing WT CDC42 set to 100.

### Statistical analyses

Statistical analyses were performed using the GraphPad Prism 10.0.2 software. Results are shown as means +/-SEM and the significance levels were calculated using one-way ANOVA (*: P<0,05; **: P<0,01; ***: P<0,001; ****: P<0,0001).

## Author contributions

JD designed and supervised the project. JD and IM provided funding. AI and RT conducted the experiments. REM performed the structural characterization. SD, GP, MN-I, RTAvW, FB, AAdJR, SC, PLAvD, AI, RG-M, TY, MT and IM provided patients’ samples. AI and JD wrote the first draft of the manuscript with contributions from SD and IM. All authors edited the paper.

## Acknowledgments

AI is supported by an international PhD fellowship from *Université Paris Cité* and a European Society for Immunodeficiencies fellowship. SD is supported by the Research Foundation – Flanders (FWO Grant number 11F4421N). IM is a senior clinical investigator at FWO Vlaanderen (supported by a KU Leuven C1 Grant C16/18/007, by FWO Grant G0B5120N and by the Jeffrey Modell Foundation). JD is supported by Inserm. This project was supported by Inserm, *Centre National de la Recherche Scientifique*, *Université Paris Cité*, *Agence Nationale de la Recherche* (2019, RIDES), AIRC (IG 28768) and Italian Ministry of Health (5×1000).

We thank: the patients and healthy donors for their participation in this study; Didier Busso (CIGEx facility), Sébastien Jacques and Franck Letourneur (GENOM’IC facility), Souganya Many (CYBIO facility) for technical assistance; Thomas Henry, Serge Bénichou and Marie-Louise Frémond for sharing plasmids and cells; Nadège Bercovici, Clotilde Randriamampita, Fabienne Régnier and Paolo Pierobon for helpful discussions.

## Supplemental material

### CLINICAL CHARACTERIZATION OF CDC42 PATIENTS

#### Patient 1

P1, carrying the R186C CDC42 mutation described in (5), was a girl born from healthy unrelated parents with no history of genetic disease. At birth, she presented with high fever, a diffuse erythematous skin rash, hepatosplenomegaly and failure to thrive. Blood analysis showed elevated inflammatory markers. A bone marrow (BM) biopsy, performed at the onset of the disease, revealed fibrosis with dyshematopoiesis. Treatment with glucocorticoids and daily therapy with anakinra improved the fever and rash but had no effect on the cytopenia. Therefore, treatment with G-CSF was started with partial response. Following tapering and/or discontinuation of glucocorticoids, a recurrence of inflammatory symptoms was observed. At 11 months, she had several episodes of intestinal bleeding. At 2 years and 6 months, she presented with three episodes of generalized seizures. Cerebral MRI was suggestive for central nervous system (CNS) inflammation in the absence of infection. The episodes were treated with high doses of glucocorticoids with good response. From the age of 5 years old, she developed four episodes of Hemophagocytic Lymphohistiocytosis (HLH). All episodes, except the final one, were resolved with treatment with high dose glucocorticoids and cyclosporine-A. During the last episode, she did not respond to high-dose glucocorticoids and IL-1 inhibition, and finally required surgical resection and consequent ileocolostomy due to massive intestinal ischemia and necrosis. However, the addition of emapalumab, an anti–IFN-γ antibody, induced a rapid resolution of the episode.

Serial measurements of the IFNα-2a concentration in the plasma of this patient was performed during the course of the disease. She displayed a high level of IFNα-2a at the onset of disease, which decreased under therapy with glucocorticoids and anakinra (Figure 6), as demonstrated in time point 1. At time point 2, P1 had an adenovirus and rhinovirus infection. Despite continuous treatment with glucocorticoids, anakinra and cyclosporine, the IFNα levels remained high. The patient finally underwent a Hematopoietic Stem Cell Transplantation (HSCT), which restored normal IFNα-2a concentration (point 4).

#### Patient 2

P2, who carried the R186C CDC42 variant and described in (5), was a male born from healthy unrelated parents. He presented at birth with persistent fever, skin rash, hepatosplenomegaly, failure to thrive, increase in inflammatory markers and pancytopenia, requiring red blood cell and platelet transfusions. Dyshematopoiesis and some lymphohistiocytic aggregates, without significant hemophagocytosis, were discovered by BM biopsy. The disease course was characterized by a persistent inflammatory state despite treatment with glucocorticoids, high doses of immunoglobulins and cyclosporine-A. Suspecting an autoinflammatory condition, treatment with anakinra was also started with only partial improvement of the clinical and laboratory parameters. At 7 months, he developed a severe HLH with multiorgan failure and rapidly progressed to death.

IFNα levels were measured at different time points in the plasma during the course of the disease. At the onset of disease, the IFNα-2a concentration was very high (Figure 6) and decreased after the addition of Anakinra, as indicated in time point 1.

#### Patient 3

P3, carrying the R186C CDC42 mutation and described in (8), was the first male child born from non-consanguineous parents. Neonatally, he presented with fever, erythema, hepatosplenomegaly and pancytopenia. He did not demonstrate dysmorphic features.

Treatment with high-dose immunoglobulins was only partially effective and addition of glucocorticoids was necessary to stabilize the patient’s condition. Tapering of the glucocorticoids induced a recurrence of fever and erythema. Since re-escalation of prednisolone could not suppress inflammation, Etanercept was added to the treatment. The inflammation could not be controlled and the patient died at 4.5 months due to overwhelming inflammation.

IFNɑ-2a levels were measured at 2 different time points in the patient’s serum. The C-reactive protein (CRP) level was low at the first time point (1.7 mg/L) and elevated at the second time point (15.6 mg/L). At the onset of disease, the IFNα-2a concentration was high and decreased after the initiation of the treatment with Etanercept, as indicated in time point 1 (Figure 6).

#### Patient 4

P4 who carries the R186C CDC42 variant was described in (10). She is a girl who presented at birth with a rash, grade II brain hemorrhage, hepatosplenomegaly and thrombocytopenia. She had frequent febrile episodes and chronic cytopenia. Her anemia and thrombocytopenia were transfusion-dependent. The parents were non-consanguineous. There was a poor response to glucocorticosteroids and rituximab. At the age of 5 months, she started on anakinra with improvement of the rash and fever. Although the dose of Anakinra was gradually augmented, there was an amelioration of the anemia, thrombocytopenia, hyperferritinemia and systemic inflammation but no complete resolution. Upon switch of Anakinra to Canakinumab, an improvement of the anemia and thrombocytopenia was noticed and the white blood cell count normalized. Yet she still suffered from recurrent episodes of anterior cervical lymphadenitis and chronic hepatosplenomegaly. The patient is currently 10.5 years old and receiving treatment with canakinumab (Figure 6).

#### Patient 5

P5, carrying the *192C*24 mutation in CDC42, is a 15-year-old male patient described in (10). His parents are non-consanguineous. He presented neonatally with fever, erythematous rash, hepatosplenomegaly, anemia and thrombocytopenia. At 7 weeks, he had an intracranial hemorrhage secondary to the severe thrombocytopenia. Until the age of 8 months, transfusion-dependent anemia and thrombocytopenia, hepatosplenomegaly, recurrent fever and rash persisted. Glucocorticoids and anakinra were started at the age of 8 months, based upon a clinical suspicion of NOMID, which resulted in a resolution of the systemic inflammation and cytopenias. He suffered from respiratory syncytial virus–induced pneumonia, septic arthritis, and cervical lymphadenitis. Recurrent infections resolved upon discontinuation of the steroids. Currently, the patient’s systemic inflammation characterized by fever, rash, anemia, and thrombocytopenia has resolved and there is an absence of hepatosplenomegaly. The patient is currently 15 years-old and receiving treatment with canakinumab (Figure 6).

#### Patient 6

P6, who carried the *192C*24 CDC42 mutation, was described in (9). She presented in early childhood with a short stature resembling achondroplasia, subtle dysmorphic features, hepatosplenomegaly, subglottic stenosis, scleritis, and inflammation of both auricles. During childhood, she suffered from recurrent infections including multiple pneumonias and an episode of viral meningitis. Blood analysis revealed elevated inflammatory markers, mild anemia, slightly elevated liver enzymes and an isolated IgM deficiency. Hypercellularity with trilineage hematopoiesis interpreted as reactive changes was discovered by bone marrow biopsy. She suffered from several pneumonias which necessitated hospitalization and intravenous antibiotic therapy. Due to the clinical suspicion of relapsing polychrondritis and an underlying primary immunodeficiency, she was treated with glucocorticosteroids and IgG maintenance therapy. When she was 44, a partial jejunum resection was performed after she suffered from acute abdominal pain due to intestinal necrotizing vasculitis. At age 55, she was admitted with acute abdominal pain and was diagnosed with a paralytic ileus. During admission, her clinical status rapidly declined and eventually she succumbed due to hemodynamic instability and pulmonary hypertension (Figure 6).

#### Patient 7

P7, carrying the Y64C CDC42 variant and described in (16), was born from non-consanguineous parents. She was small for gestational age, microcephalic, hypotonic and mildly dysmorphic. She manifested psychomotor developmental delay, feeding difficulties and growth retardation. She had microretrognathia, hypertelorism with incomplete eyelid closure, depressed nasal bridge, strabismus, severe astigmatism, arachnodactyly, and sensorineural hearing loss. She developed scoliosis, pes planus, camptodactyly, fingernail exostosis, hepatosplenomegaly with multiple spleen and kidney hyperechogenic lesions (which were not biopsied), liver hemangiomas, and a thoracic-abdominal aortic aneurysm. She suffered recurrent lower respiratory tract infections since childhood with a diagnosis of bronchiectasis at age 15 years despite antibiotic treatment and physiotherapy. Laboratory evaluations at that time showed profound lymphopenia, especially of B cells, neutropenia, anemia, and thrombocytopenia. A bone marrow biopsy at age 16 showed dysmegakaryopoiesis but otherwise normocellular marrow with normal differentiation. She had hypogammaglobulinemia in the first year of life with IgA and IgM deficiency, but only IgA deficiency persisted later in life. At age 25 years, she experienced weight loss (> 10%), increased coughing, fever, and elevated inflammatory markers. C-reactive protein (CRP) and erythrocyte sedimentation rate (ESR) increased. HLH could not be confirmed. The patient’s state did not improve despite broad antifungal and antibiotic treatment, but responded to steroids. Bone marrow biopsy showed myeloid hypercellularity, erythroid hypoplasia, and fibrosis grade 2/3 according to the WHO classification of myeloid neoplasm, suggestive of primary myelofibrosis. As part of the hematology workup, a NGS panel (Illumina platform) on peripheral blood was performed to screen for driver variants in *JAK2*, *CALR*, and *MPL*, as well as for non-driver variants in a large set of genes. No somatic or germline variants could be identified. At age 26 years, she was admitted with cachexia, dyspnea, and hypoxemia. A chest CT scan showed bilateral ground glass opacities, honeycombing, and a crazy paving pattern with cyst formation. Bronchus’ aspirate revealed MRSA and HSV-1 for which she was treated accordingly. Ventilatory support was escalated to extracorporeal membrane oxygenation (ECMO) from D+7. Suspecting an inflammatory component, she was started on 80 mg methylprednisolone per day on D+11, after which she could be weaned from ECMO. Upon tapering of steroids, lung infiltrates flared—a pattern seen twice, after which glucocorticoid therapy was maintained from D+41. During this course, she had several episodes of pulmonary hemorrhages. CT angiography showed several hypertrophic bronchial arteriae which were embolized. At D+41 after admission, *Stenotrophomonas maltophilia* was cultured from the sputum. Ultimately, the patient succumbed to shock associated with *Stenotrophomonas maltophilia* bacteremia at D+68 after admission.

---

**Supplementary Table 1:**
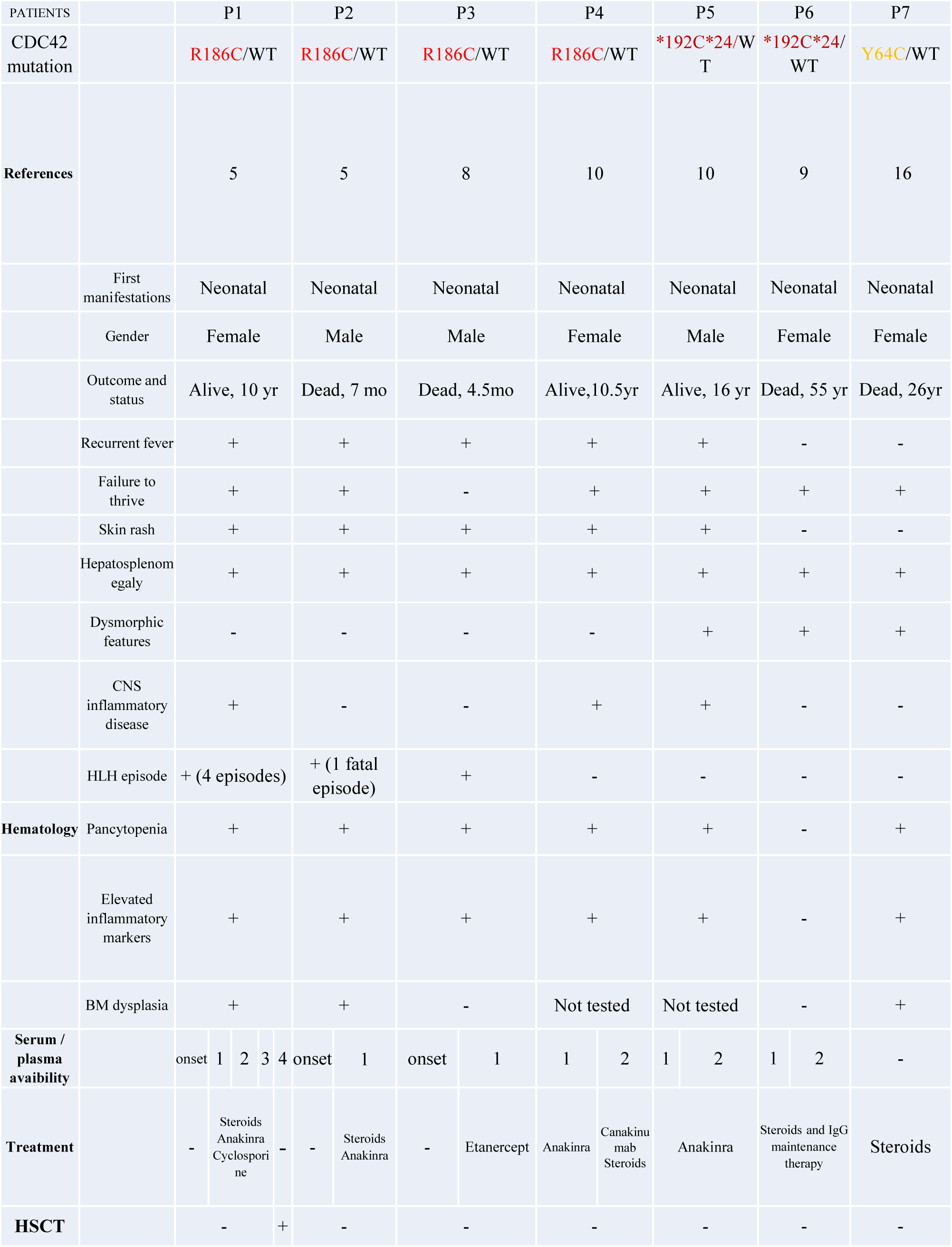
Clinical features of the CDC42 patients studied here. CNS: central nervous system. HSCT: Hematopoietic Stem Cell Transplantation

**Supplementary Table 2:**
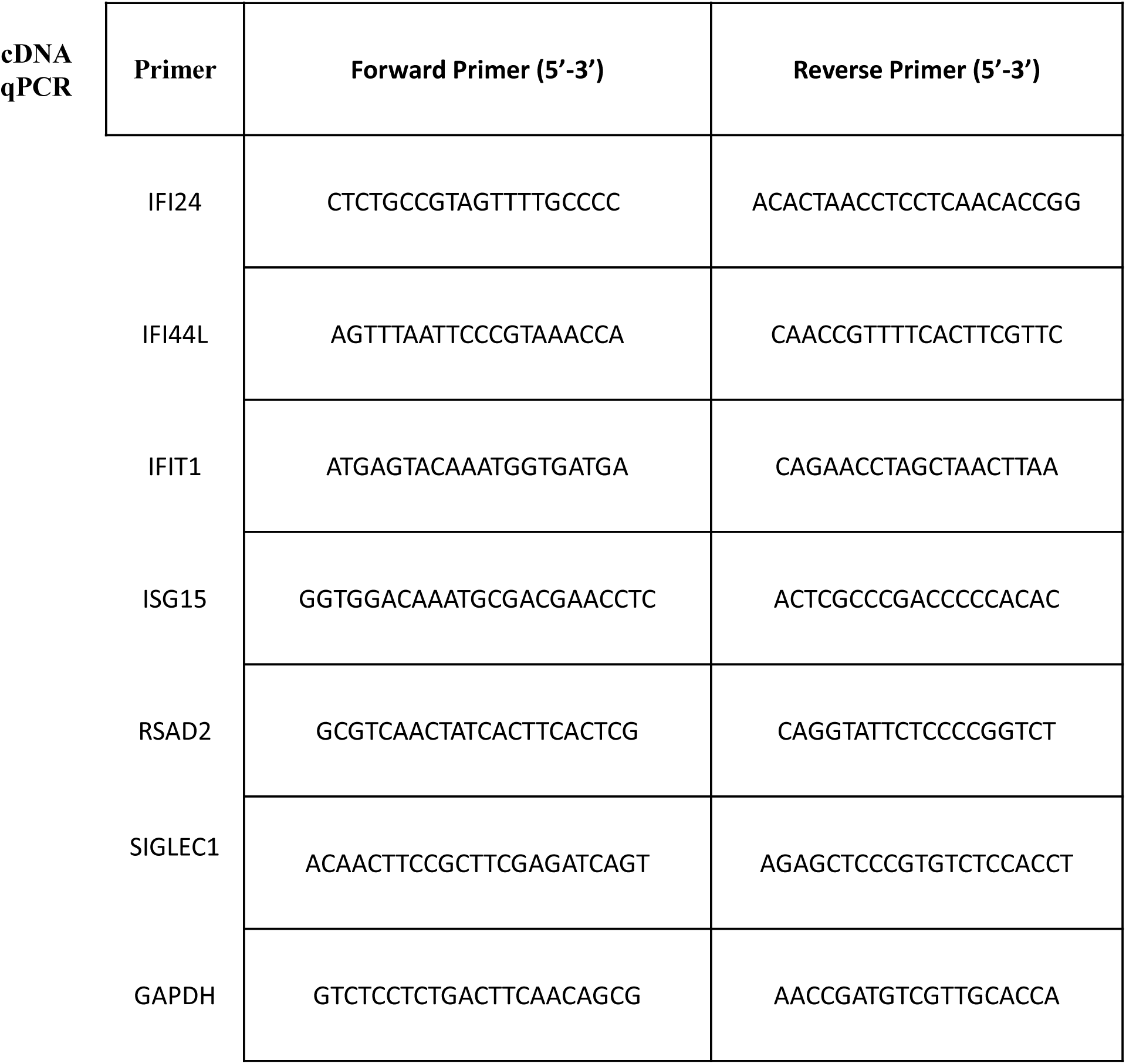
qPCR primers used in this study.

**Supplementary Table 3:** List of staining reagents used here.

| Primary Antibody | Host species | Source | Catalog Reference |
| --- | --- | --- | --- |
| Anti-PC-1 | Rabbit polyclonal | Rockland | 600-401-103s |
| Anti-GM130 | Mouse monoclonal | BD Biosciences | 610822 |
| Anti-SERCA2 ATPase | Goat polyclonal | Abcam | Ab219173 |
| Anti-Giantin | Rabbit Polyclonal | Abcam | ab80864 |
| Human STING-TMEM173 | Mouse monoclonal | R&D System | #723505 |
| Anti-STING (D2P2F) | Rabbit monoclonal | Cell Signaling | 13647s |
| Anti-P-STING (SER366) | Rabbit | Cell Signaling | #19781 |
| Anti-P-STAT1 (Tyr 701) | Rabbit monoclonal | Cell Signaling | CST7649 |
| Anti-P-IRF3 (Ser 396) | Rabbit Polyclonal | Thermo fisher | 720012 |
| Anti-GRP78 BiP | Rabbit Polyclonal | Abcam | ab21685 |
| Phalloidin-647 |  | Thermo fisher | A22287 |
| Hoechst |  | Thermo fisher | H21486 |
| Donkey anti Mouse IgG- Alexa 647 |  | Invitrogen | A32787 |
| Donkey anti Rabbit IgG- Alexa 647 |  | Invitrogen | A31573 |
| Donkey anti Rabbit IgG- Alexa 568 |  | invitrogen | A10042 |
| Donkey anti Goat IgG- Alexa 488 |  | Invitrogen | A11055 |
| Donkey anti Rabbit IgG- Alexa 488 |  | Invitrogen | A21206 |

